# DTI-CDF: a CDF model towards the prediction of DTIs based on hybrid features

**DOI:** 10.1101/657973

**Authors:** Yan-Yi Chu, Yu-Fang Zhang, Wei Wang, Xian-Geng Wang, Xiao-Qi Shan, Yi Xiong, Dong-Qing Wei

**Affiliations:** State Key Laboratory of Microbial Metabolism, School of Life Sciences and Biotechnology, Shanghai Jiao Tong University, Shanghai, China; School of Mathematical Sciences, Shanghai Jiao Tong University, Shanghai, China

**Keywords:** Drug-target interaction prediction, cascade deep forest, hybrid features

## Abstract

Drug-target interactions play a crucial role in target-based drug discovery and exploitation. Computational prediction of DTIs has become a popular alternative strategy to the experimental methods for identification of DTIs of which are both time and resource consuming. However, the performances of the current DTIs prediction approaches suffer from a problem of low precision and high false positive rate. In this study, we aimed to develop a novel DTIs prediction method, named DTI-CDF, for improving the prediction precision based on a cascade deep forest model which integrates hybrid features, including multiple similarity-based features extracted from the heterogeneous graph, fingerprints of drugs, and evolution information of target protein sequences. In the experiments, we built five replicates of 10 fold cross-validations under three different experimental settings of data sets, namely, corresponding DTIs values of certain drugs (*S*_*D*_), targets (*S*_*T*_), or drug-target pairs (*S*_*P*_) in the training set are missed, but existed in the test set. The experimental results show that our proposed approach DTI-CDF achieved significantly higher performance than the state-of-the-art methods.

## 1 Introduction

Drug discovery is the process of identifying new candidate compounds with potential therapeutic effects, during which the prediction of drug-target interactions (DTIs) is an essential step [1, 2]. Drugs are significant in the human body by interacting with various targets. Proteins represent an important type of target, and their functions can be enhanced or inhibited by drugs to achieve phenotypic effects for therapeutic purposes [3, 4]. However, the number of drug candidates approved by the FDA is relatively small [4, 5], mainly due to the possible adverse effects of multitargeting of drugs. Currently, large quantities of researches have focused on DTIs prediction because it is an essential tool in the context of drug repurposing. Since the high cost of experimental determination of DTIs, it is thus necessary to develop efficient computational methods by making use of the heterogeneous biological data on known DTIs to understand the mechanisms of action of drugs in the human body.

The recent computational approaches are mainly concerned with the ligand-based approaches [6], docking simulation approaches [7], and chemogenomic approaches [8]. The ligand-based interactions are motivated by the idea that similar molecules generally bind to similar proteins. However, these methods perform poorly especially when the number of known ligands is insufficient. The docking simulation methods need three-dimensional structure information of proteins for simulation, thus being intractable when numerous proteins are not access to their structure information. The chemogenomic methods have attracted much interest because it maps both the chemical feature space and the genomic feature space into a unified Euclidean space, namely pharmacological space and obtains good performance in DTIs prediction. These methods can mainly be divided into three classes: graph-based method, network-based method, and machine learning-based methods [9]. The former two methods sometimes have encountered the dilemma for the scarcity of known DTIs and unidentified negative DTIs data, thus the results are not highly confidential. However, machine learning-based methods can better overcome this dilemma for its advanced capability to sufficiently exploiting the sample information, resulting in more reliable prediction results than the former two methods. On the other hand, there are large quantities of high efficient machine learning methods have been employed in various research fields and obtain good performances, which motivates us to make full use of these methods such that enhances the prediction accuracy. Moreover, the machine learning models are being studied intensively which thus provides broaden possibilities for the improvement of DTIs prediction.

The recent machine learning-based methods are composed of semi-supervised models and supervised models. As for semi-supervised machine learning method, Xia et al. [10] first designed a manifold Laplacian regularized the least square (LapRLS) by using the concept of the bipartite local model (BLM) [11] with the assist of labeled and unlabeled data. Subsequently, some improved LapRLS-based models were used for DTIs prediction, such as NetLapRLS [10] and ILRLS [12]. In addition, there are some interesting work, such as the Network-Consistency-based Prediction method [13] carried out by maximizing the known interaction’s rank coherence, the PUDT method [14] predicted DTIs by using positive unlabeled data, a set of Graph Auto-Encoder-based models applying multi-view similarities [15], and the NormMulInf method [16] developed a principal component analysis model based collaborative filtering method by using low-rank similarity matrix. These methods are easy to be implemented. However, these methods are unable to apply to drugs without any targets information and more time-consuming as the result of the unlabeled negative interactions and thus cannot be implemented on a large-scale database.

In the context of supervised learning of which known interactions are labeled positive samples and other interactions are labeled negative ones, there are two categories methods: similarity-based methods and feature vector-based methods. As for the similarity-based methods[17], the key assumption is the “guilt-by-association”, i.e., similar drugs tend to share similar targets and vice versa. Based on the nearest neighbor method, a variety of similarity measures such as Jaccard similarity, Cosine similarity, and Pearson correlation similarity are exploited to calculate the similarity score such that enhances the model prediction ability. Shi et at. [18]designed a Similarity-Rank-based predictor to present the likelihood of each drug-target pair tending to interact or not. The merit of this method does not need the complex parameter optimization. Chen et al. [19] employed an inference model based on the random walk on the heterogeneous network. Cheng et al. [20] developed three inferring networks consisting of drug-based similarity inference network, target-based similarity inference network and network-based inference to discover DTIs. The method cannot predict new drug or targets. Based on the BLM, the DTIs prediction problem can be transformed into a binary classification problem. Thus, various classifiers have been exploited such as regularized least square classifier [21], support vector machine (SVM) classifier [22]. Mei et al. [23] further exploited BLM with Neighbor-based Interaction Profile Inferring, which adds a preprocessing component to infer training data from neighbors’ interaction profiles. In addition, in the matrix factorization methods that are typically utilized in recommendation systems to find a potential user-item relationship, DTIs can be transformed into a matrix completion problem aiming to find missing interactions. Gönen et al. [24] presented a pioneer work of Kernelized Bayesian Matrix Factorization with Twin Kernels. There are other methods such as Probabilistic Matrix Factorization [25], multiple Similarities Collaborative Matrix Factorization [26], Graph regularized matrix factorization [27], weighted GRMF [27] and Variational Bayesian Multiple Kernel Logistic Matrix Factorization [28]. These methods improve the interpretability to some extent by using the knowledge of matrix theory, however, are not practical in large-scale samples for the high cost of the matrix completion problem. Based on the feature vector, a variety of methods have been proposed. Wang et al. [29] employed a restricted Boltzmann machine to find the data distribution such that identifies DTIs relationship as well as drug modes of action. Based on meta-path-based topological features, Fu et al. [30] employed random forest classifier to do prediction, and Zong et al. [31] calculated it with SkipGram model. These methods are difficult to find new targets or drugs in known networks. Olayan et al. [32] exploited a novel method called DDR to solve the above problem and improve the prediction accuracy which executes graph mining technique firstly to acquire the comprehensive feature vectors and then applies the random forest model by using different graph-based features extracted from the drug-target heterogeneous graph. Farshid Rayhan et al. [33] explored sampling method to avoid data imbalanced problem and used Adaboost model to do prediction, and it is the first time combine evolutionary and structural information of proteins as a part of the feature vector. Ezzat et al. [34] carried out the ensembling learning method which using Decision Tree and Kernel Ridge Regression as base classifiers. Besides, Wen et al. [35] proposed a deep learning framework by deep belief network for the first time applying in this field, which needs further exploration. Based deep learning method, Hu et al. [36] used Auto-Encoders to learn representations as SVM’s feature vector, and Tian et al. [37] used a deep neural network to learn and predict.

Motivated by the previous studies [38, 39], we propose a cascade deep forest (CDF) model that further improves the performance of predicting DTIs. This method combines the above two machine learning-based methods. First, we utilize FP2 fingerprint (FP2), to extract the structural information of drugs, Pseudo-position specific scoring matrix (PsePSSM) to extract evolution information of protein sequence, and adds Path-category-based multi-similarities feature (PathCS) based on the heterogeneous graph of DTIs. Then, we apply the CDF model under three experimental settings through five repeated 10-fold cross-validations in four representative data sets, and the performance evaluation is performed using both AUPR and AUC metrics. Besides, the statistical hypothesis test is used to evaluate the results’ significance. Finally, we verify that the proposed DTI-CDF method is significantly better than the current state-of-art methods available.

## 2 Materials and Methods

### 2.1 Data sets

This study uses four data sets compiled by Yamanishi et al. [40] to evaluate the performance of the proposed DTI-CDF method in DTIs prediction. The four data sets are distinguished and named by the target protein of the drug: enzymes (E), ion channels (IC), G-protein-couples receptors (GPCR) and nuclear receptors (NR). These data sets contain known human DTIs retrieved from the KEGG BRITE [41], BRENDA [42], SuperTarget[43] and DrugBank [44] databases. Therefore, it is generally considered as the gold-standard data sets.

In order to simulate more practically, in these four data sets, we consider the entire space of the DTIs, where the number of known DTIs (or positive samples) is much lower than the number of unknown DTIs (or negative samples). Thus, these four data sets are very unbalanced, as shown in Table 1.

**Table 1.** Summary of quantitative information for the four data sets

| Data sets | Known interactions | Unknown interactions | Drugs | Targets |
| --- | --- | --- | --- | --- |
| NR | 90 (6.41%) | 1314 (93.59%) | 54 | 26 |
| GPCR | 635 (3%) | 20550 (97%) | 223 | 95 |
| IC | 1476 (3.45%) | 41364 (96.55%) | 210 | 204 |
| E | 2926 (1%) | 292554 (99%) | 445 | 664 |

### 2.2 Feature construction

#### 2.2.1 PathCS

PathCS [32] is based on the heterogeneous DTIs weighted graph, containing drugs, targets, and their similarities or interactions. In this graph, the edge between two target nodes or two drug nodes represents their similarities and the weight *w*_*x*_ is their similarity value. The edge between a target and a drug denotes a known drug-target interaction and the weight is equal to 1.

There are six kernels used in this study to generate similarity profiles for drugs and targets which have been proved a more robust and less redundant similarity set [32], their information as follows:

1. Protein kernels. We use the proteins’ amino acid sequence information to generate the spectrum kernel [45] and set the subsequence length *k* as 4.
2. Drug kernels. There are three side-effects kernels as drug information sources. The first resource obtained from the SIDER database [46], which contains information on marketed drugs and their adverse reactions. For each side-effect classification, a binary (absence or presence) profile was used to represent drugs. The other two pharmacological profiles are derived from the FDA’s adverse event reporting system [47] based on the frequency and binary information, respectively, of side-effects classifications. These three pharmacological profiles are used to generate similarity profiles through the weighted cosine correlation coefficient. And if a drug is not in the data resources, its assigned similarity is 0.
3. Gaussian Interaction Profile kernel (GIP kernel). The GIP kernel [21] is a binary matrix based on the DTIs network for drugs or targets, in which the absence or presence of interaction in the network for each drug or each target is described as 0 or 1, respectively. However, this kernel cannot be computed for a new drug (or a new target), which do not have any drugs (or targets) to interact with the training data set. To solve this problem, we adopt the method of neighbor-based interaction-profile inferring [23] to calculate this kernel.

After obtaining the above similarity measures, the first step is to combine the drugs’(or targets’) multiple similarity measures into one fused matrix [48] to build a heterogeneous DTIs graph, then extract PathCS for each drug-target pair. The path category is defined by a path structure that starts at a drug node and ends up at a target node such as to set the path length to 2 or 3. Path categories are as follows: drug-drug-target, drug-target-target, drug-drug-drug-target, drug-drug-target-target, drug-target-drug-target and drug-target-target-target. We define two normalized matrices 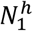 and 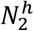 according to the above 6 path categories *C*^*h*^, *h* = 1,2,…, 6. For a specific drug *d*_*i*_ and a specific target *t*_*j*_, we denote one path from *d*_*i*_ to *t*_*j*_ as *p*_*q*_ and the set of paths is *R*_*ijh*_. In addition, the path between *d*_*i*_ and *t*_*j*_ is built by the intermediate nodes which are restricted to be the 5 nearest neighbors of *d*_*i*_ and *t*_*j*_, respectively. Thus, the 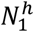 and 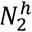 with elements 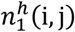 and 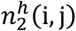, respectively, are computed as follows:

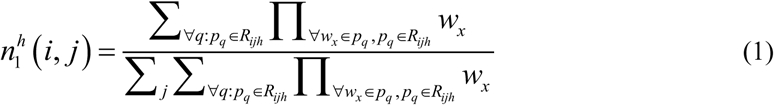

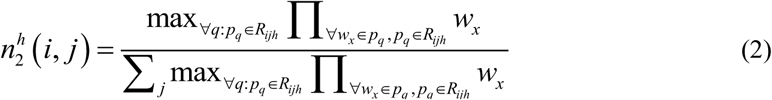

#### 2.2.2 PsePSSM

There are varieties of methods that can be used to extract characteristic information on proteins based on the amino acid sequence. In this study, we select PsePSSM [49] as our protein description method. This method combines the position-specific scoring matrix (PSSM) [50] and the pseudo amino acid composition (PseACC) [51], which represent evolution information and sequence information, respectively. For an amino acid sequence A with L residues, the dimension of a normalized PSSM is L × 20 as follows:

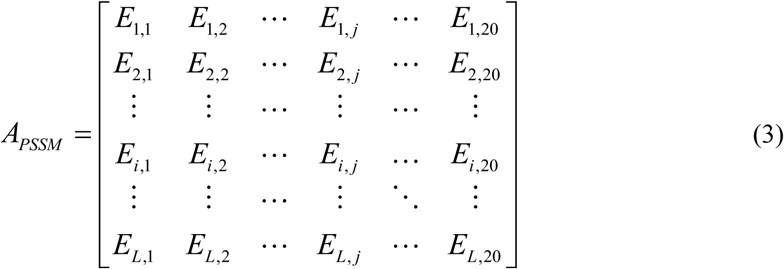

where *i* is the position of the residue in the amino acid sequence, *j* is the type of one of the 20 native amino acids, *E*_*i,j*_ is the score at which the *i*-th residue in the amino acid sequence is mutated to the *j*-th amino acid.

For proteins with different amino-acid sequence length, the number of rows of this matrix is different. To solve this problem, one can turn this matrix into a vector of length 20 as

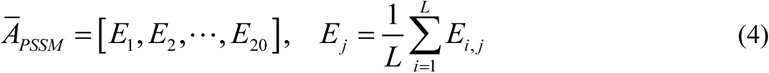

where *E*_*j*_ is the average occurrence score of all residues in the amino-acid sequence that is mutated to the j-th amino acid during evolution. The PseACC is used to describe an amino-acid sequence A using Eq. (3) as

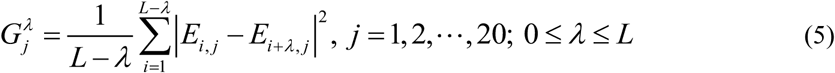

where 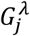 is the λ-tier correlation factor of the jth amino acid and λ denotes the difference order along each column of the matrix *A*_*PSSM*_. For clearly, 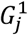 represents the relevant factor calculated by the 1-order difference of row elements along the jth column of the matrix, i.e., the closest PSSM score, on the protein sequence of amino acid type *j*, and 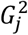 represents the 2-order difference along the *j-*th column of the matrix, namely the second closest PSSM score and so on. In this study, λ ∈ {1,2,… 10} because of its best performance [49], thus generating a vector containing 200 components. Hence, a protein using the PsePSSM method obtains a feature vector with 220 elements, namely 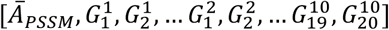.

#### 2.2.3 FP2

FP2 [52] is a path-based fingerprint to express drugs. This fingerprint identifies all ring and linear substructures of lengths from 1 to 7 in the molecule, among them, the C and N substructures should be excluded. Then, these substructures are mapped to a 1024-bit string. Thus, for each molecule, FP2 is a hexadecimal 256-dimensional vector.

### 2.3 Classification algorithm

Firstly, we generate the feature vector for each DTI. The GIP similarity is constructed according to training data, then PathCS is obtained. Based on the drug-target pair, the PathCS, FP2, and PsePSSM are merged to form input feature vector, called hybrid features.

Secondly, CDF classifier [53] is used to predict DTIs. The input of the CDF model is hybrid features. Then, the new category probability vector link input feature vector is used as the next layer input, and the final category probability vector is output through multiple learners. When building a CDF model (Fig.1), it is important to determine the machine learner used for each layer. In the model, we set the number of learners of each layer from 2 to 6, and Random Forest (RF) [54] and XGBoost (XGB) [55] are used as learners to follow the “good but distinguishable” principle. In addition, the depth of layers is identified automatic by the trend of evaluation metrics.

**Fig. 1.**
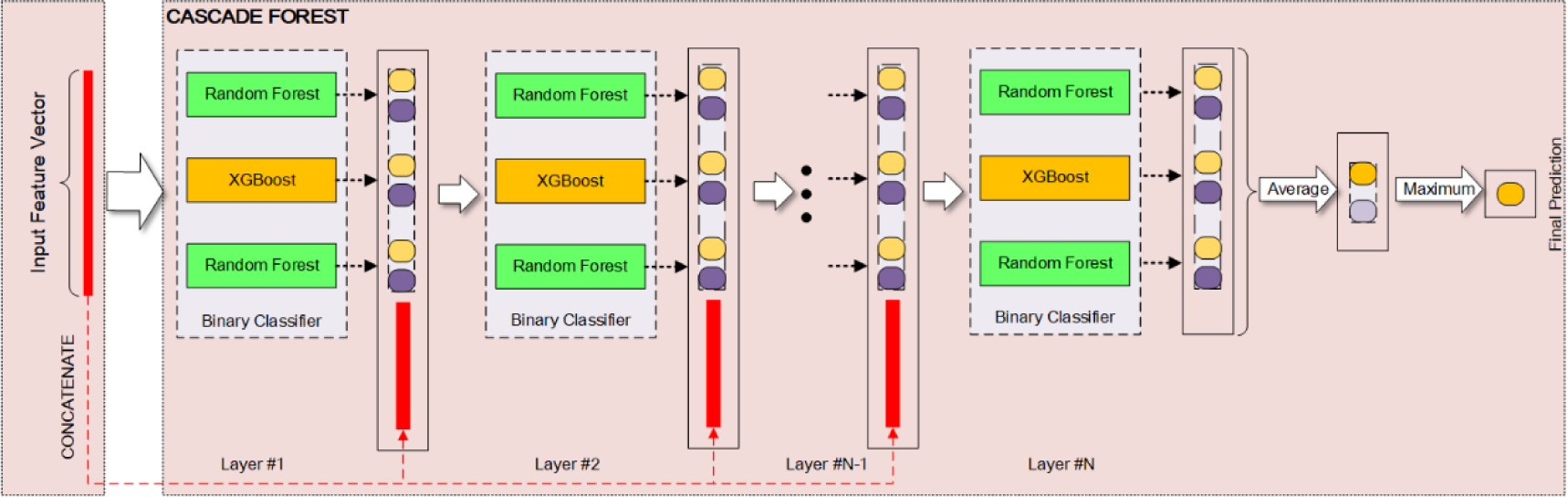
This machine learning model is composed of an input feature vector, a CDF classifier, and a final prediction. In particular, CDF is the core unit of the model, which has five variants in this study.

In each variant, each layer consists of a different number of RF and XGB binary classifiers and different layers own the same structure. The figure shows one special model in which each layer has two RF learners and one XGB learner, denoted as RF2-XGB1. Other variants are RF2, RF3, RF3-XGB2, RF4-XGB2, respectively.

### 2.4 Experimental settings and cross-validation

In this study, we evaluate three experimental settings as Table 2 shows, which includes most of the conditions for DTIs prediction. In Table 2, objects are new indicates that no corresponding DTIs in the training data, and known vice versa. In order to facilitate the comparison with other methods, we followed previous studies [32, 56-58] as the benchmark and conducted the 10-fold cross-validations (CVs) test for each experimental setting of each data set, and the above process was repeated 5 times using different random seeds.

**Table 2.**
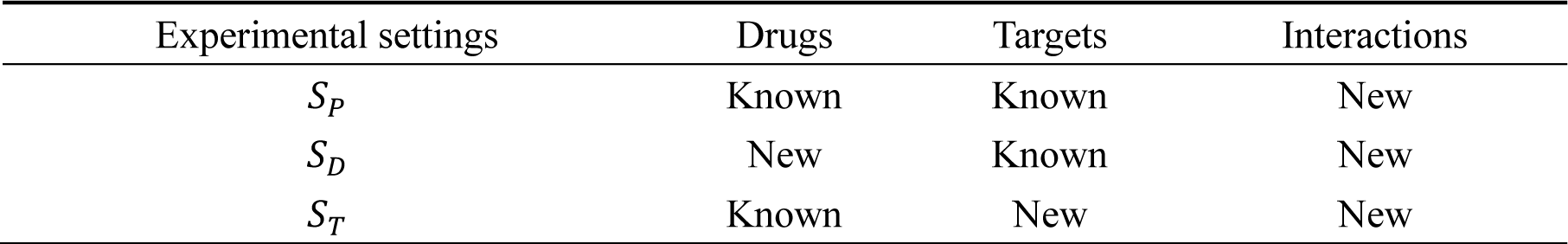
Summary the corresponding DTIs information in test data of three experimental settings

| Experimental settings | Drugs | Targets | Interactions |
| --- | --- | --- | --- |
| $S_P$ | Known | Known | New |
| $S_D$ | New | Known | New |
| $S_T$ | Known | New | New |

### 2.5 Performance evaluation

For each fold of each predictive model, the following metrics are calculated:

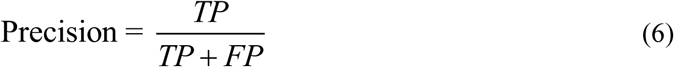

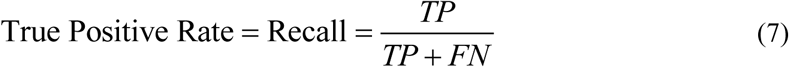

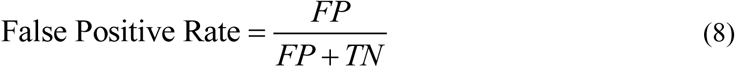

where *TP* is true positive, *FP* is false positive, *FN* is false negative and *TN* is true negative. We plot the precision-recall curve (PR curve) based on different precision and recall, and the receiver operating characteristic curve (ROC curve) based on different recall and false positive rate, respectively, under the condition of different classified cutoff values. We define AUPR and AUC as the area under the PR curve and the ROC curve, respectively. For each experiment setting of each data set, the AUPR and AUC are calculated as a measure of model performance as follows:

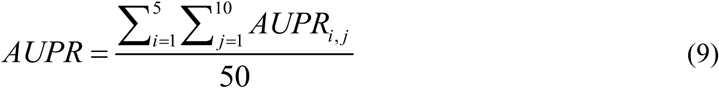

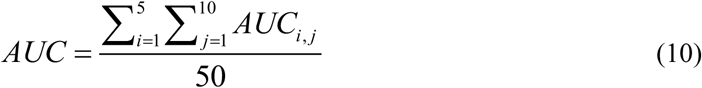

where *i* represents the *i*-th repeated trials, *j* represents the *j*-th fold of CVs. Since the positive samples and negative samples in each data set are highly unbalanced, the AUPR provides a better performance estimate relative to AUC because it more severely penalizes the false positives[59].

## 3 Results and Discussion

### 3.1 Predictive ability of different types of features

The aim of this section is to evaluate the effect of adding FP2 and PsePSSM into a similarity-based feature vector. For computational method in DTI prediction, extracting the features of drugs and targets effectively is very important. In previous studies, there are two main methods for generating features of drugs and targets: (i) based on the chemical structure for drugs and the amino acid sequence for proteins to extract features. For chemical structures, various molecular fingerprints of compounds can be used, such as FP2 [49], Extended-Connectivity Fingerprints [60], etc. The amino acid sequences can be represented by amino acid composition [61], PseAAC [62], PsePSSM [49], etc. (ii) based on association rules, drugs with similar chemical structures tend to bind similar proteins. It is based on heterogeneous networks of DTIs, using single or fusion similarity measures as features [23, 31, 32, 58].

The similarity-based feature plays a crucial role in predicting DTIs. Large quantities of studies use evolution information for targets’ sequence and structure information for drugs, PsePSSM and FP2 are reported that they can effectively extract information on drugs and targets, respectively, as good results are obtained by using the above two features [32, 49]. In this study, we combined the above two feature extraction method to generate the input feature vector. Based on PathCS, we constructed hybrid features by adding FP2 and PsePSSM to describe drugs and targets, respectively. The results show that this hybrid features achieves higher AUPR and AUC than using PathCS along (Fig. 2). Furthermore, it demonstrates that in the prediction of DTIs, combining the structure information of drugs and evolution information of targets’ sequence with similarity information could increase accuracy than similarity information used alone.

**Fig. 2.**
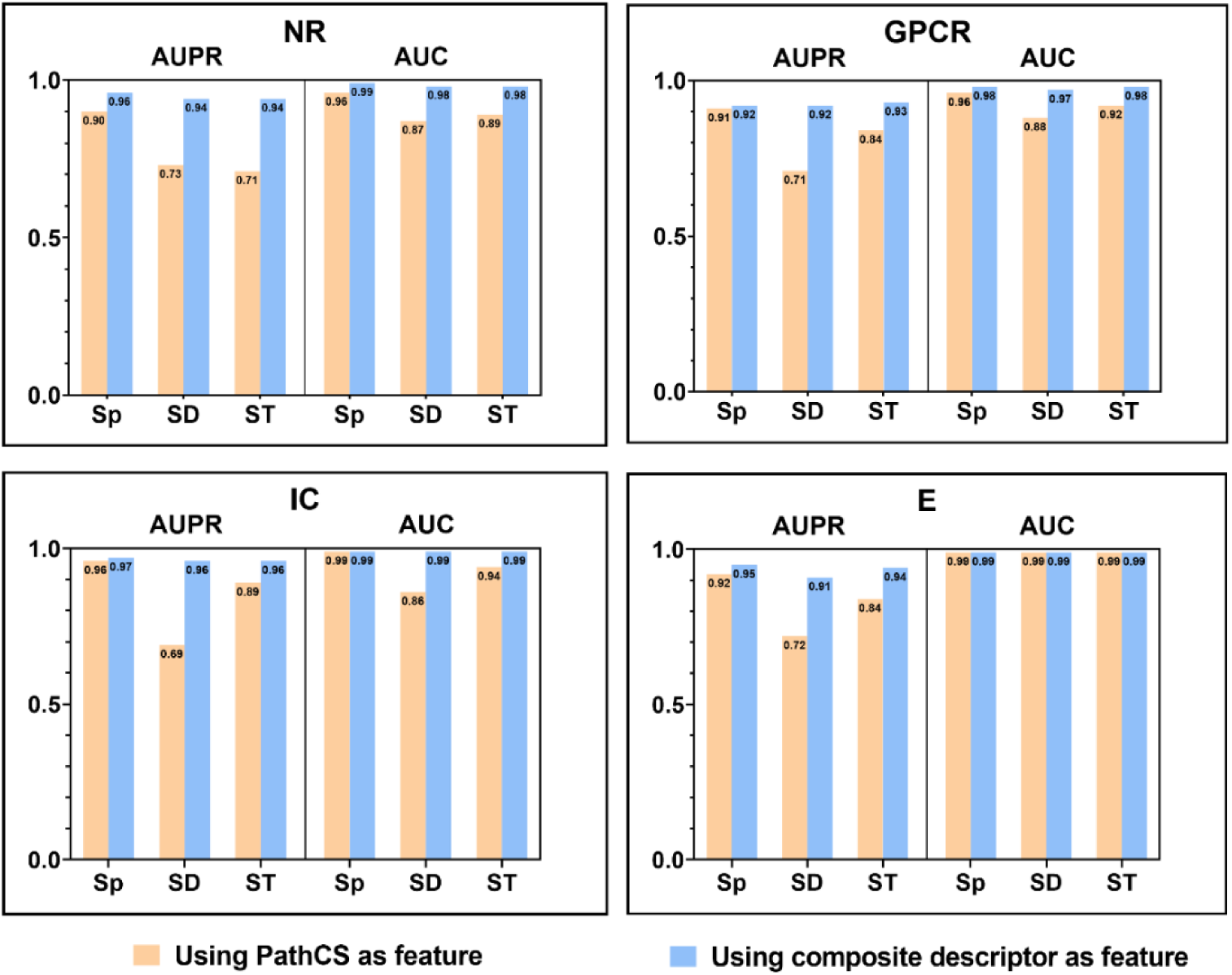
Comparison of DTI-CDF methods that using PathCS only and hybrid features as a feature vector

In addition, we use a statistical hypothesis test [63] to further explore the extent to which hybrid features outperform single PathCS used in the DTI-CDF method. Differences in the results of different prediction methods are caused by a variety of factors, such as data composition, training model and experimental setting, etc. In order to exclude other factors and only consider the difference caused by the prediction method, the one-sided paired t-test, that is a pairwise comparison method based on paired data, is employed. Firstly, the difference *d*_*i*_ ∈ **D**, (i = 1,2,…, 24) of AUPR and AUC based on 12 experimental conditions (i.e. four data sets under three experimental settings) between the above two methods are calculated. It is assumed that the difference *d*_*i*_ are all from the normal distribution *N*(*μ*_*d*_, *σ*^2^), where both *μ*_*d*_ and *σ*^2^ are unknown. Then, a statistical hypothesis test is performed based on the data obtained above. If the hybrid features used in DTI-CDF method is not different from the single PathCS, the difference *d*_*i*_ between each pair of data belongs to a random error, and the random error can be considered to obey a normal distribution with a mean of zero. Assuming that there is no difference between the above two methods, the test hypothesis is as follows:

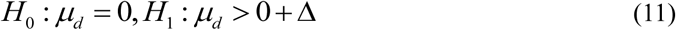

By the *t*-test of a single population means using the normal distribution, the rejection domain is:

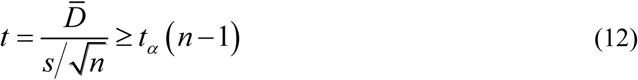

where 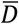 is the mean of the sample, *s* is the standard deviation of the sample, *n* is the sample size, *α* is the significance level and Δ is equivalent to the effect size of mean difference, defined as 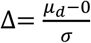. In order to ensure that only when the hybrid features are far superior to the single PathCS used in DTI-CDF method, it can be tested with a high probability 1 − β, we set *α* = 0.05, Δ= 0.9, *β* = 0.01. Under these conditions, a sample size *n* not less than 21 is required, and the actual sample size *n* is 24 satisfying the requirement. The rejection domain and the actual effect size of mean difference are

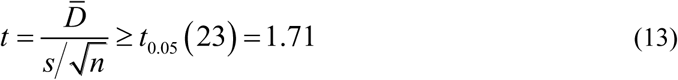

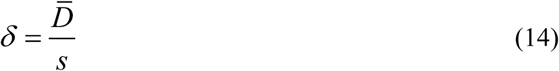

Substituting *d*_*i*_ into the above formula yields the observed value *t*_0_ of *t*, then the *p*-value of the right-tailed *t*-test can be calculated by

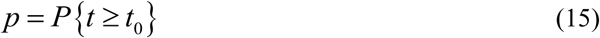

The calculation results show in Table 3, illustrated that at the significance level *α* = 0.05, *t* falls within the rejection domain, so the null hypothesis *H*_0_ is rejected and the alternative hypothesis *H*_1_ is accepted. By calculation, it is known that the minimum significance level *p*-value of rejecting the null hypothesis *H*_0_ is far less than *α*, and the effect size of the mean difference *δ* more than 0.8. We can reasonably believe that when training the model, using the hybrid features as the input feature vector is significantly better than just applying PathCS.

**Table 3.** The hypothesis test results

| | $t_0$ | $p$-value | $\delta$ |
| --- | --- | --- | --- |
| Compare hybrid features with PathCS. | 6.11 | $1.56 \times 10^{-6}$ | 1.25 |
| Compare CDF model with RF model. | 2.03 | 0.03 | 0.41 |
| Compare DTI-CDF method with the DDR method. | 6.11 | $1.57 \times 10^{-6}$ | 1.25 |

In order to explore the reason why the combination of the three types of features improves the performance of the model, we analyzed the correlation between these three types of features. We performed a cross-correlation function analysis on the three types of features used in the model training on 12 experimental conditions. The cross-correlation function *ρ*_*xy*_ of two vectors *x* and *y* is defined as:

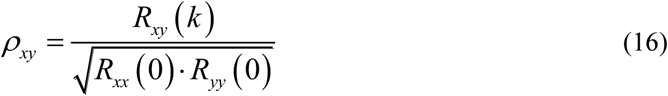

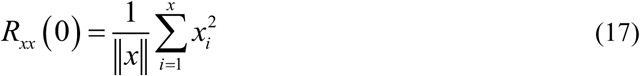

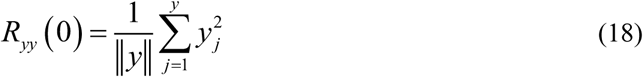

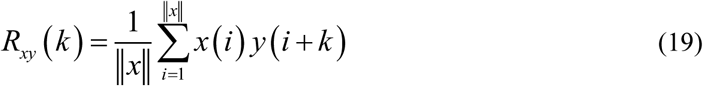

for *k* = 0,1,2…, ‖*y*‖ − ‖*x*‖ + 1, where ‖*x*‖ is the length of *x*, ‖*y*‖ is the length of *y*, and assume ‖*y*‖ ≥ ‖*x*‖. For each experimental condition, the data is first separated into three parts according to the three types of features and flattened separately to obtain three corresponding vectors with different lengths, of *N* × *M*, where *N* is the number of the samples and *M* is the dimension of a feature vector. Then, the cross-correlation function among the three types of features is calculated according to Eq. (17-20). According to the characteristics of the flattened data, we make 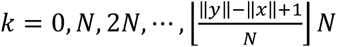 to reduce the computational cost. We choose the maximum value of the absolute value from *ρ*_*xy*_ as the correlation coefficient to measure the linear correlation between two types of features to ensure the reliability of the results, this value is in the range of [0,1], equal to 0 and 1 represent linear independent and complete linear correlation, respectively. Since the correlation coefficients of three experimental settings on a particular data set are very close, we selected the maximum value shown in Fig. 3. It can be seen that under the four data sets, the correlation coefficients among the three types of features are all below 10^−4^, indicating that the data of different features are very close to irrelevant, thus, these three types of features can be considered to be orthogonal to each other, which can produce the greatest information when describing DTIs. Therefore, after adding FP2 and PsePSSM into a feature vector, the representation of DTIs is enriched without redundancy, thereby avoiding over-fitting and improving the performance of the model.

**Fig. 3.**
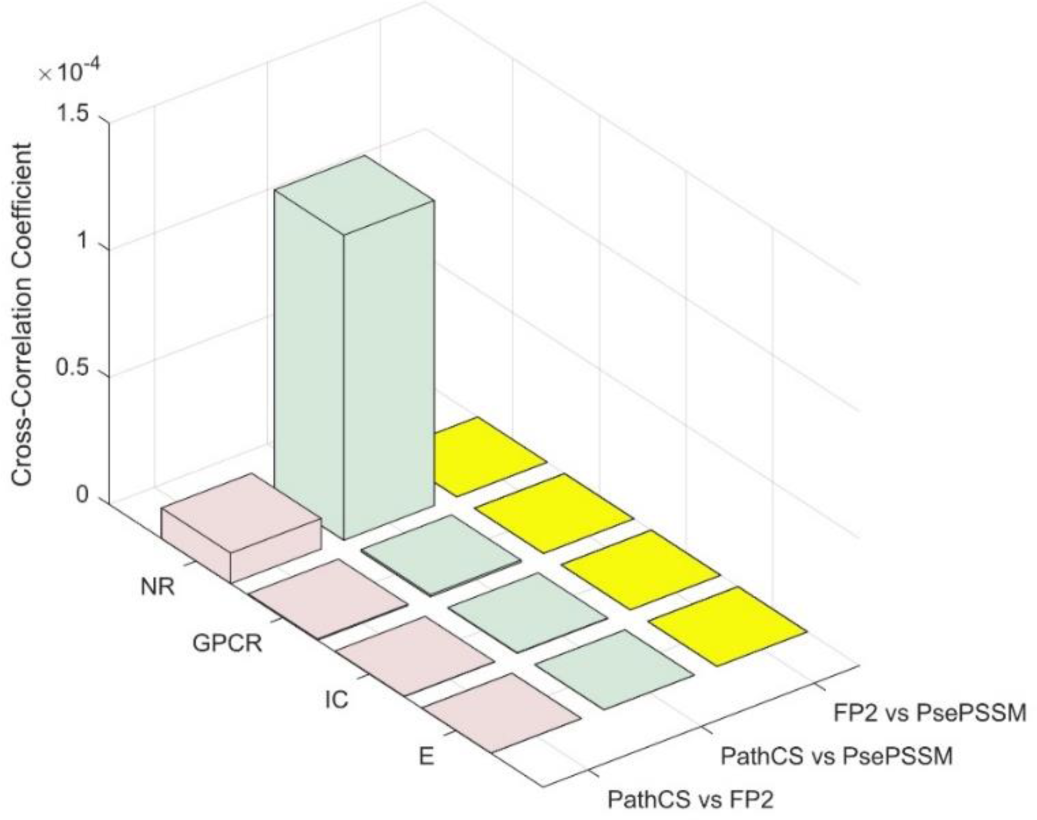
Comparison of cross-correlation coefficients among three types of features

### 3.2 Effect of CDF model on the DTI-CDF performance

In the CDF model, one or more RF or XGB learners are added based on a single RF learner. Compared to a single RF, we demonstrate that the performance of CDF is better at predicting DTIs. In this regard, we used PathCS as input feature vector to train CDF model, then compared with the DDR method through AUPR and AUC metrics, as the DDR method uses PathCS to train the single RF model. Note that the experimental conditions of the above two methods are identical, such as the division of data sets, random state, etc. We observed that in the CDF model, the results are higher than those in the RF model except that the AUPR of the IC and E data sets under the experimental setting *S*_*D*_ are approximately equal to that of the RF model (as shown in Fig. 4). In the comparison of AUC, it was found that half of the results showed that the CDF model was higher than the RF model. In the other half, except for the IC data set under the experimental setting *S*_*D*_, the difference between other results is no more than 3%. It is worth noting that for highly unbalanced data sets, AUPR is more accurate than AUC. Therefore, we have reason to think that the CDF model is better than the RF model. In addition, we prove this statistically with the hypothesis test.

**Fig. 4.**
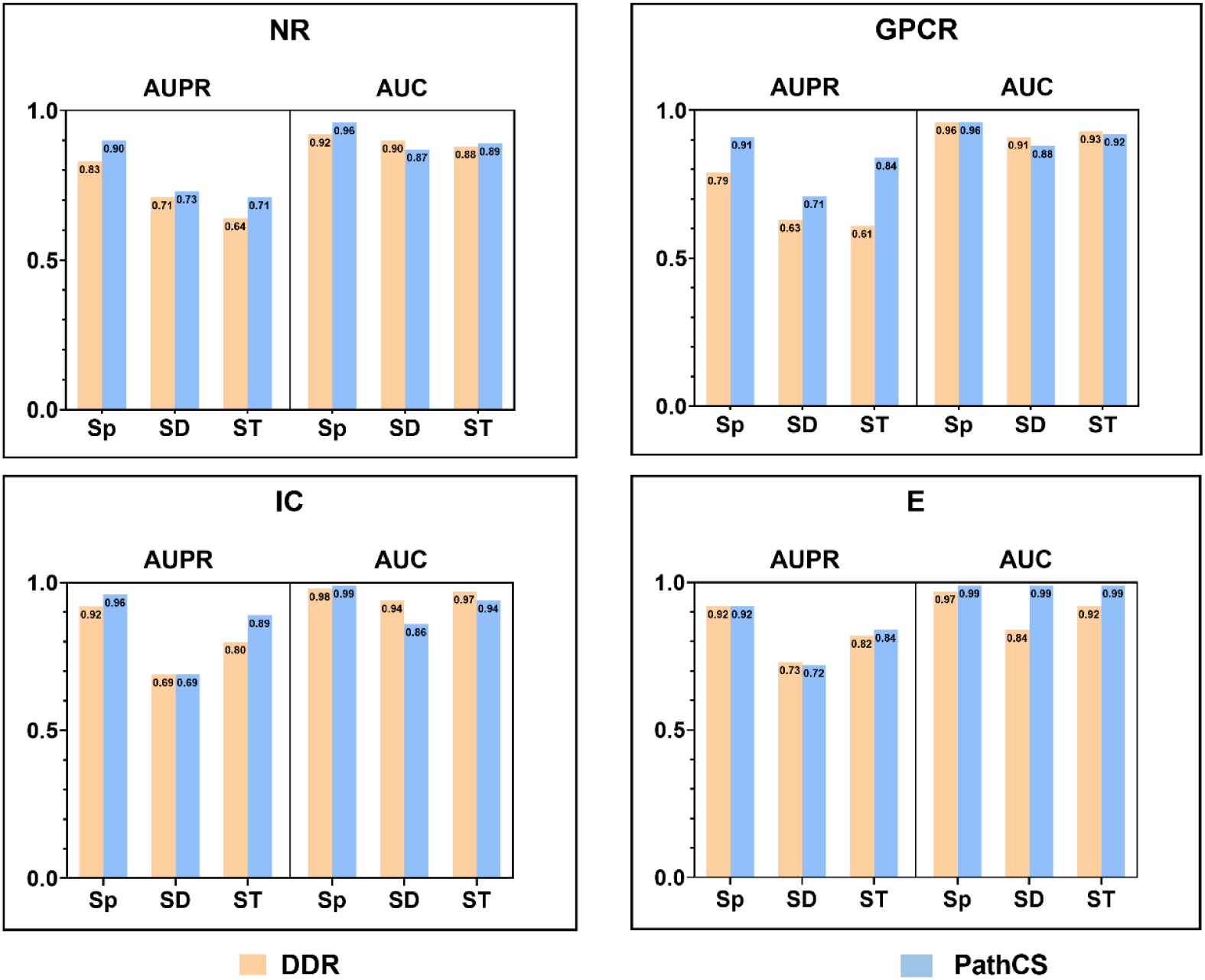
Comparison of DTI-CDF method that using PathCS as a feature vector and DDR method

Firstly, the difference between the above two methods of AUPR and AUC is calculated. Assuming that there is no significant difference between these two methods, the statistical hypothesis is still expressed by Eq. (12). According to t-test, the corresponding rejection domain is Eq. (14). The test results are shown in Table 3.

The calculation results show that at the significance level *α* = 0.05, *t* falls within the rejection domain, so the null hypothesis *H*_0_ is rejected and the alternative hypothesis *H*_1_ is accepted. By calculation, it is known that the minimum significance level *p*-value of rejecting the null hypothesis *H*_0_ is less than *α*, and the effect size of mean difference *δ* is 0.41. We can reasonably believe that using CDF model is medium significantly better than using the RF model for DTIs prediction.

It is not difficult to find that all the results under the experimental setting *S*_*p*_ are very good (i.e. both AUC and AUPR are more than 90%) when using the CDF model, while the results of the *S*_*T*_ and *S*_*D*_ experimental settings are not, a similar phenomenon also appears in the results of the DDR method. The reason is that the model has been over-fitting under the experimental settings of *S*_*D*_ and *S*_*T*_, because the AUPR and AUC on the training set are both more than 0.9 when using CDF model. There are many reasons for over-fitting, the most important reason is that the model is too complex or the feature is not sufficient to describe the sample. From the above results, we can see that although the CDF model introduced by us is more complex than the RF model, it still achieves better results. Therefore, the crux of the over-fitting problem lies in the feature, which will be echoed in Section 3.1 and Section 3.3.

### 3.3 Comparisons with the state-of-the-art algorithms (CDF vs. DDR)

Based on these four data sets, the DDR method [32] is proved as the most advanced method for predicting DTIs under the same experimental conditions (i.e. 5 repeated trails of 10-fold CVs under three experimental settings of each data set), so we only compare the proposed DTI-CDF method with the DDR method. Experiments show that DTI-CDF achieves better performance than DDR under the same conditions (Fig. 5), and all AUPRs exceeds 0.90, all AUCs are more than 0.96.

**Fig. 5.**
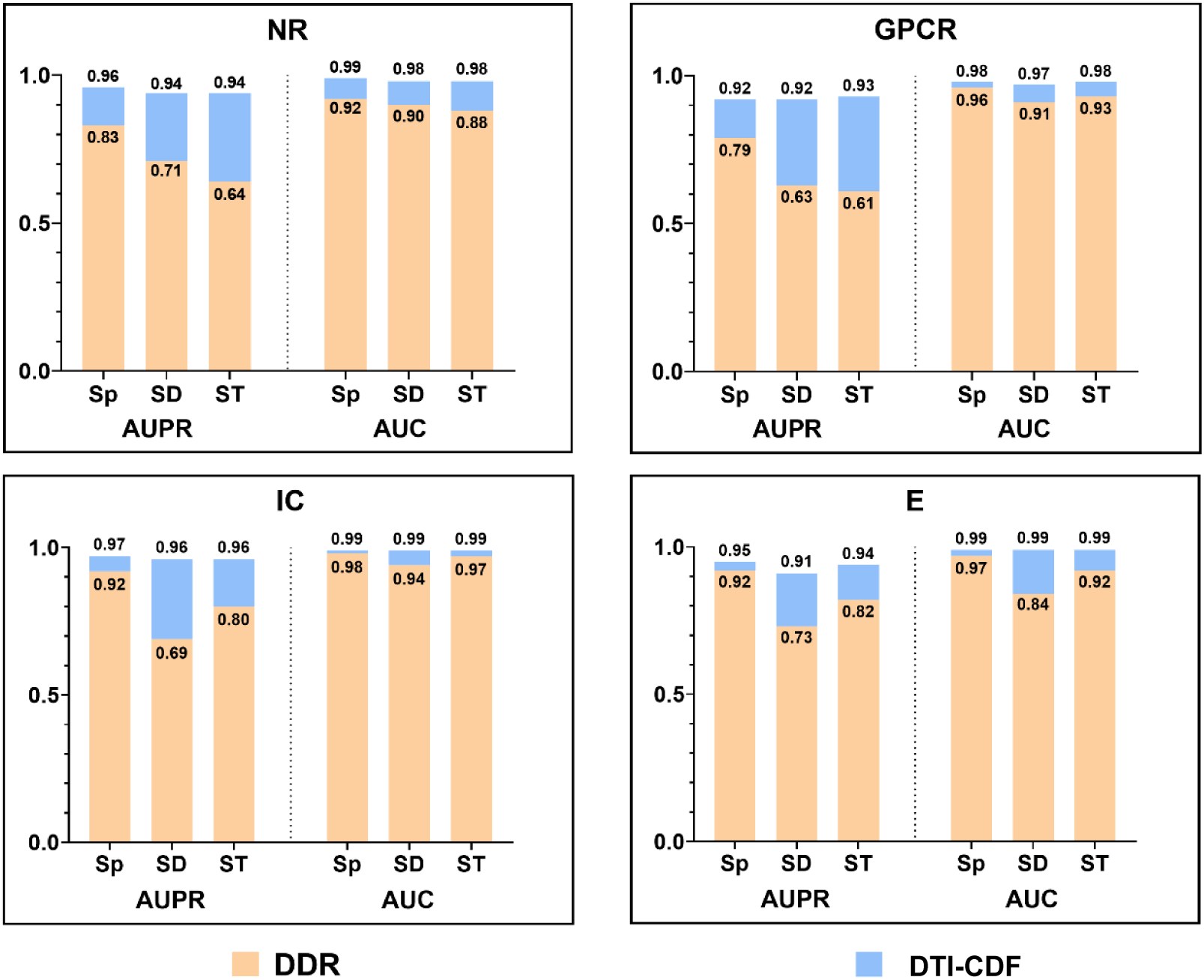
Comparison of the AUPR and AUC of DTI-CDF with the DDR methods.

In order to compare the degree of difference between the DTI-CDF method and the DDR methods, we also carried out the one-sided paired t-test. Similarly, the AUPR and AUC of two methods under the same experimental conditions are subtracted in pairs. Assuming that there is no significant difference between the DTI-CDF method and the DDR method, Eq. (12) and Eq. (14) are still used to the test hypothesis and the rejection domain. The test results are listed in Table 3. The calculation results show that at the significance level *α* = 0.05, *t* falls within the rejection domain, so the null hypothesis *H*_0_ is rejected and the alternative hypothesis *H*_1_ is accepted. By calculation, it is known that the minimum significance level *p*-value of rejecting the null hypothesis *H*_0_ is far less than *α*, and the effect size of mean difference *δ* is larger than 0.8. We can reasonably believe that the DTI-CDF method is highly significantly better than the DDR method.

## 4 Conclusions

We propose a DTI-CDF method to predict DTIs, which utilizes similarity information for drugs and targets, structural information for drugs, and evolution information for targets’ sequence to obtain feature vector as the input of CDF algorithm for DTIs prediction. We use AUPR and AUC to evaluate the performance of the DTI-CDF method under three different experimental settings based on gold-standard data sets, all of them more than 0.9 and superior than the current state-of-art DDR method. It is further proved that the performance of the DTI-CDF method is significantly better than other existing methods when a known DTI is missing from the training data, especially in searching targets for new drugs (*S*_*D*_ setting) and finding drugs for new targets (*S*_*T*_ setting). This demonstrates that the DTI-CDF method has a higher predictive ability for the real scene of DTIs statistically. In addition, we have demonstrated that combining the similarity, structural and sequence information as feature vectors can better describe DTIs. We firstly use CDF algorithm in this field and prove its superiority. In the future, we plan to use the DTI-CDF method to deal with the regression problem such as calculating the affinity between drugs and targets.

